# Evaluating HIV-1 Infectivity and Virion Maturation Across Varied Producer Cells with a Novel FRET-Based Detection and Quantification Assay

**DOI:** 10.1101/2023.12.25.573317

**Authors:** Aidan McGraw, Grace Hillmer, Jeongpill Choi, Kedhar Narayan, Dacia Marquez, Hasset Tibebe, Taisuke Izumi

## Abstract

The maturation of HIV-1 virions is a crucial process in viral replication. Although T cells are a primary source of virus production, much of our understanding of virion maturation comes from studies using the HEK293T human embryonic kidney cell line. Notably, there is a lack of comparative analyses between T cells and HEK293T cells in terms of virion maturation efficiency in existing literature. We previously developed an advanced virion visualization system based on the FRET principle, enabling the effective distinction between immature and mature virions via fluorescence microscopy. In this study, we utilized pseudotyped, single-round infectious viruses tagged with FRET labels (HIV-1 Gag-iFRETΔEnv) derived from Jurkat (a human T lymphocyte cell line) and HEK293T cells to evaluate their virion maturation rates. HEK293T-derived virions demonstrated a maturity rate of 81.79%, consistent with other studies and our previous findings. However, virions originating from Jurkat cells demonstrated a significantly reduced maturation rate of 68.67% (p < 0.0001). Correspondingly, viruses produced from Jurkat cells exhibited significantly reduced infectivity compared to those derived from HEK293T cells, with the relative infectivity measured at 65.3%. This finding is consistent with the observed relative maturation rate of viruses produced by Jurkat cells. These findings suggest that initiation of virion maturation directly correlates with viral infectivity. Our observation highlights the dynamic nature of virus-host interactions and their implications for virion production and infectivity.

## Introduction

Human Immunodeficiency Virus-1 (HIV-1) continues to pose a significant global health challenge. The advancement of combination antivirals has dramatically improved the management of HIV-1, targeting various stages of the virus lifecycle, including entry, reverse transcription, integration, and protease activity [1-6]. Recently, the development of capsid inhibitors has emerged as a promising strategy for long-acting antiviral treatments against HIV-1 [7-12]. These novel inhibitors, targeting critical steps in both the early and late stages of the virus lifecycle, notably disrupt the formation of infectious particles. This innovation underscores the need for a comprehensive understanding of the virion maturation process, a key phase in the HIV-1 life cycle where non-infectious particles are transformed into infectious virions. Our previous research developed a unique tool for detecting virus maturation utilizing a fluorescence resonance energy transfer (FRET)-based technique [13]. This approach enables the semi-automatic evaluation of virion maturation through fluorescent microscopy, effectively minimizing human bias. It offers crucial insights into the structural changes that are vital for virus infectivity. Building upon this, our current study explores virion maturation in different cellular environments. Although T cells are recognized as the primary source for HIV-1 replication [14-20], most existing research assessing viral maturation predominantly utilizes the HEK293T human embryonic kidney cell line [21-24]. Utilizing the FRET-based system, we generated infectious FRET-labeled viruses in both HEK293T and Jurkat cells, an immortal human T-cell leukemia cell line. Surprisingly, we observed that viruses produced by Jurkat cells exhibited a significantly lower maturation efficiency (approximately 69%) compared to those from HEK293T cells (approximately 82%). In alignment with the observed differences in maturation rates, the infectivity of viruses from Jurkat cells was approximately half that observed in viruses from HEK293T cells. Given that CD4+ T cells are the principal producers of HIV-1, these cells have evolved multiple antiretroviral mechanisms. For example, the HIV-1 Viral Infectivity Factor (Vif) is a crucial accessory molecule for HIV-1 replication in vivo. However, it is not necessary for HEK293T cells, as these kidney-derived cells do not express the potent antiviral host factor APOBEC3G [13, 25, 26]. It is highly plausible that kidney cells, having not been exposed to lentiviruses throughout evolutionary history, exhibit greater susceptibility to these viruses compared to T cells, which have been concurrently evolving to resist viral infections. This study is at the forefront of comparing virion maturation and infectivity among cell lines from different origins, highlighting a novel approach in this field of HIV-1 research.

## Results

### Virion Maturation Efficiency in Different Producer Cells

In this study, we quantified the proportion of mature and immature virions in FRET-labeled viruses produced by HEK293T (kidney) and Jurkat (T-cell leukemia) cell lines. In each sample, we captured a total of 21 images, from which we created binary images based on the emitted and excited images of YFP (Fig. 1B). We then calculated the FRET ratio for each virion by dividing the intensity of YFP (the FRET acceptor) by the total CFP intensity, which serves as the FRET donor. After calculating the FRET ratio for each virion, based on the extracted signal intensities of the FRET donor and acceptor, we applied kernel density estimation to the histograms of FRET efficiencies. The area where the HIV-1 Gag-iFRETΔEnv curve overlapped with the HIV-1 Gag-iFRETΔPRΔEnv curve was identified as the proportion of immature virions, as per our established methodology (Fig. 1C) [13]. In our analysis, we examined 14,021 and 22,779 particles, as well as 20,457 and 23,181 particles of HIV-1 Gag-iFRETΔEnv and HIV-1 Gag-iFRETΔPRΔEnv labeled virions produced by HEK293T and Jurkat cells, respectively, across three independent experiments. The mature virion proportion in the HIV-1 Gag-iFRETΔEnv population from HEK293T cells was 81.79% ± 0.06 (Fig. 1D), consistent with our previous findings and electron microscopic counts reported in other studies [13, 21-24]. In contrast, Jurkat cells produced mature virions at a lower rate of 68.67% ± 0.04 (Fig. 1D), which is significantly different from that of HEK293T cells (*p*-value < 0.0001). Furthermore, we quantified the Mean Fluorescent Intensity (MFI) of the YFP signal in both mature and immature virions. The results confirmed that the average incorporation of fluorescent proteins, indicated by the MFI of YFP (Fig. 1E), was similar in both mature and immature virions from HEK293T and Jurkat cells (26,691.31 ± 2,733.73 vs. 23,309.51 ± 766.93 in mature virions and 24,619.37 ± 2,857.71 vs. 24,536.10 ± 2,046.71 in immature virions, respectively, and 24,619.37 ± 2,857.71 vs. 24,536.10 ± 2,036.70 in ΔPR). These findings indicate that the number of incorporated fluorescent proteins is comparable, ensuring that the FRET energy transfer efficiency is neither under nor overestimated in virions produced by either HEK293T or Jurkat cells.

**Figure 1.**
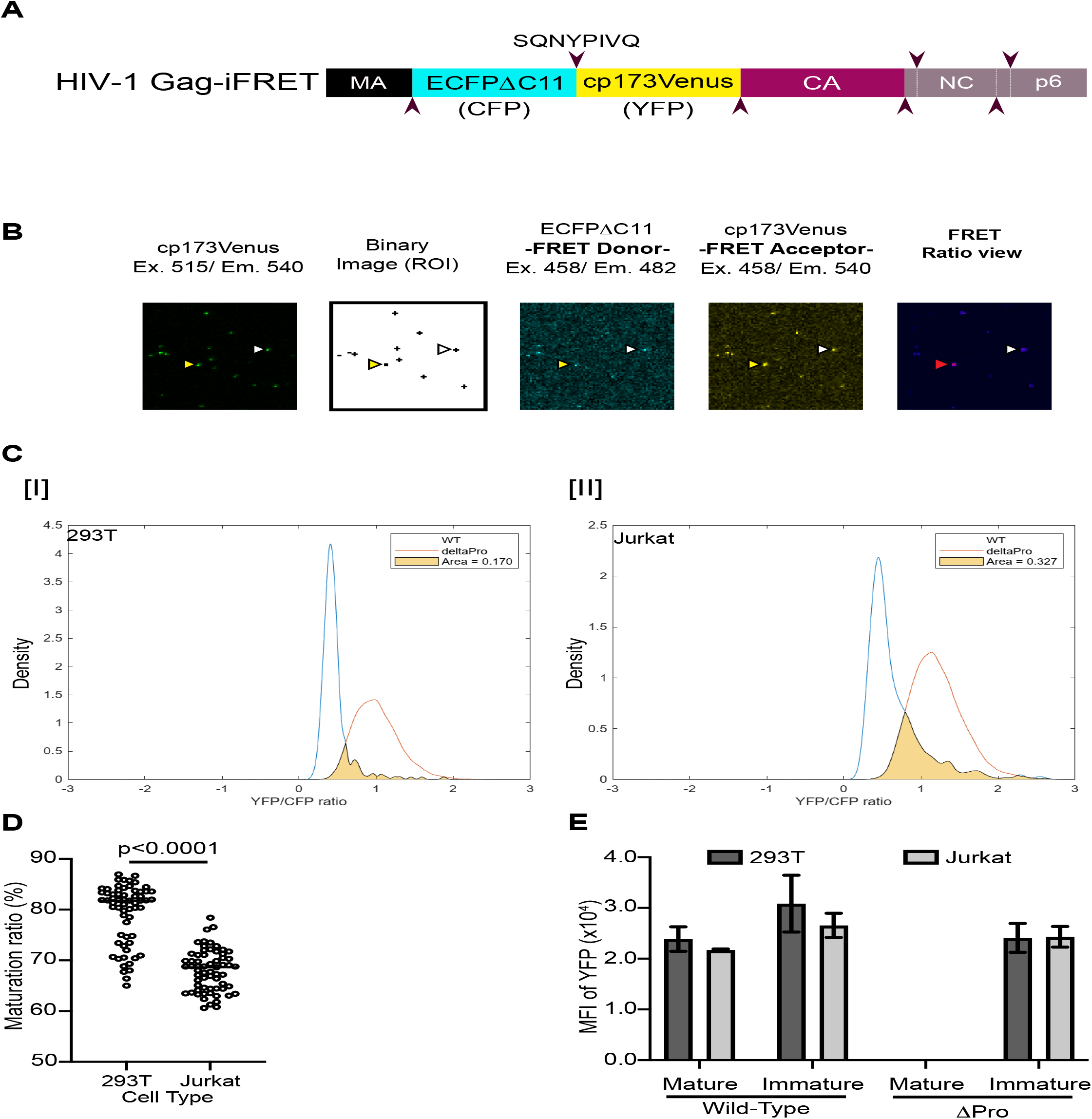
Quantitative Analysis of Mature Virions from HEK293T and Jurkat Cells. (A) Schematic representation of the HIV-1 Gag iFRET construct in the Gag region. ECFPΔC11 and cp173Venus are used for efficient single-molecule CFP and YFP FRET pairing. (B) Representative FRET-labeled HIV-1 Gag-iFRETΔEnv virion images from Jurkat cells displayed in a single field. The left panel shows an image captured through the YFP excitation (515 nm) and emission (540 nm) channels (YFP channel). The second image shows the binary image, derived from the YFP channel, identifying virus particle locations as regions of interest (ROI). The third and fourth images are captured using CFP (FRET donor) excitation (454 nm), with CFP emission (483 nm, third) and YFP emission (540 nm, fourth) channels. The right panel shows the FRET efficiency ratio view, computationally constructed from FRET donor (CFP excitation/CFP emission) and acceptor (CFP excitation/YFP emission) images. The color spectrum indicates FRET signal strength, with high FRET appearing in red (yellow arrows) and decreasing FRET shifting toward blue (white arrows). (C) Kernel Density Estimation Curves of FRET Intensity in HIV-1 Gag-iFRETΔEnv Virions Produced by HEK293T and Jurkat Cells. The curves depict the distribution histograms of FRET intensity for virions labeled with HIV-1 Gag-iFRETΔEnv from (I) HEK293T and (II) Jurkat cells. The x-axis represents the range of FRET intensity. The y-axis shows the adjusted density of curves for both HIV-1 Gag-iFRETΔEnv and HIV-1 Gag-iFRETΔPRΔEnv virions. The area under the curve for HIV-1 Gag-iFRETΔEnv overlapping with that of HIV-1 Gag-iFRETΔPRΔEnv is calculated and considered indicative of the proportion of immature virions within the total population of HIV-1 Gag-iFRETΔEnv virions. (D) Scatter Plot of Mature Virion Quantification displays the median percentage of mature virions across 21 images derived from three independent experiments, culminating in a total of 63 data points. Statistical significance was assessed using the Mann-Whitney U test. (E) Comparison of Mean Fluorescent Intensity (MFI) of YFP Signals in Mature and Immature HIV-1 Gag-iFRETΔEnv labeling Virions. The levels of fluorescent protein incorporation, as indicated by the YFP signal, were not significantly different between mature and immature virions in either the wild-type or ΔPR viruses. The error bars represent the standard error from three independent experiments. Statistical significance was assessed using the Wilcoxon matched-pairs signed rank test.

### Virus Infectivity in Different Producer Cells

To further examine the characteristic differences between viruses produced by HEK293T and Jurkat cells, we performed a single-round infection assay using TZM-bl cells to assess the infectivity of FRET-labeled viruses originating from these cell lines. The infectivity of viruses from Jurkat cells was significantly lower compared to those from HEK293T cells, measured at 65.3% ± 5.6 (Fig. 2). This discrepancy is consistent with the observed differences in the maturation rates of viruses produced by Jurkat cells. (Fig. 1).

**Figure 2.**
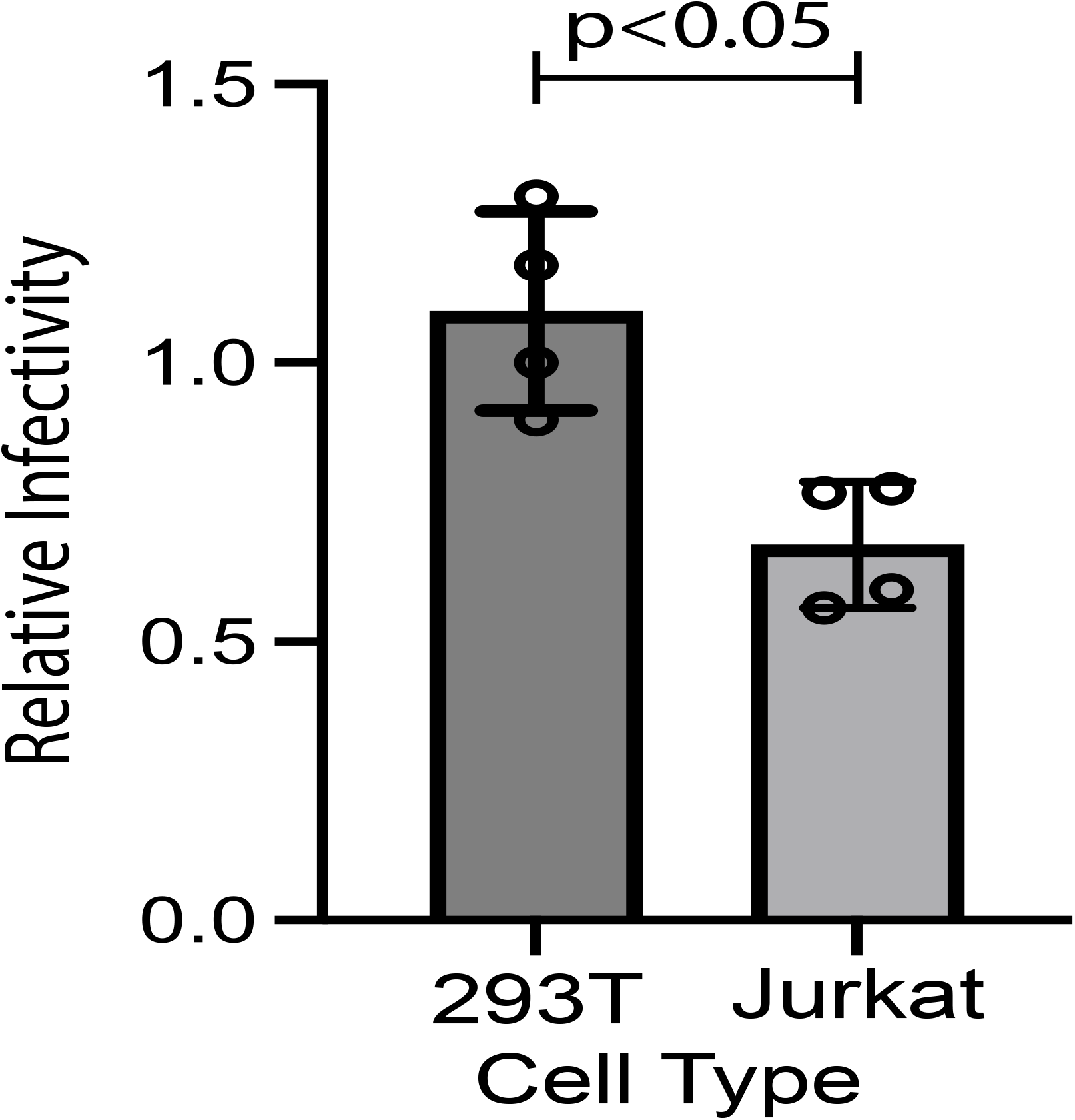
Comparing Virus Infectivity Between HEK293T and Jurkat Cells. Single-round infectivity assays using TZM-bl cells were performed to evaluate the differences in virus infectivity produced by HEK293T and Jurkat cells. The bar plot illustrates the relative infectivity of the FRET-labeled virus produced by Jurkat cells in comparison to its infectivity when produced by HEK293T cells. Error bars represent the standard deviation from four independent experiments. Statistical significance was determined using a Mann-Whitney U test.

## Discussion

The findings of this study provide groundbreaking insights into the maturation process of HIV-1 virions and their impact on viral infectivity, challenging several established concepts in the field. Our utilization of an advanced FRET-based visualization system has allowed for a detailed comparative analysis of virion maturation in two distinct cellular environments: HEK293T and Jurkat cell lines. One of the most notable findings of our study is that virus maturation and subsequent infectivity in T cell lines, which are derived from the natural targets for HIV-1, are lower compared to kidney-derived cell lines. HEK293T cells exhibited a reduced rate of immature virions (18.21%), whereas virions derived from Jurkat cells displayed a higher rate of immaturity (31.33%), correlating with its respective infectivity (65.3% ± 5.6 for Jurkat cell-derived virions compared to the infectivity observed in virions from HEK293T cells). Recent studies have provided deeper insights into the role of inositol hexakisphosphate (IP6) in HIV-1 capsid assembly, which is a critical factor in virus infectivity [27]. These studies reveal that the immature Gag lattice of HIV-1 virions enriches IP6, facilitating capsid maturation. This process is characterized by proteolysis of the Gag polyprotein by the viral protease, which induces vital conformational changes in the capsid protein (CA). These changes led to the formation of a conical, cone-shaped mature capsid consisting of over 1,000 CA copies and forming approximately 200 hexamers and 12 pentamers, collectively known as capsomers. Sowd et al. showed that IP6-depleted T-cell lines produce virion particles with incompletely cleaved Gag proteins [28]. This study indicated that virions from IP6-deficient cells exhibited lower quantities of mature MA and CA cleavage products and higher proportions of MA-CA-SP1/MA-CA cleavage intermediates than virions from wild-type cells. In the context of our FRET-based assay, which specifically detects cleavage of the Gag polyprotein between MA-CA, our FRET detection system can potentially monitor the IP6-dependent efficiency of Gag polyprotein cleavage. Furthermore, they characterized virus particle morphology using electron microscopy and calculated that mature particles in another T-cell leukemia cell line, MT-4 cells, comprised 62% of the total virions, which aligns with the results from our FRET detection system in Jurkat cells (Fig. 1C [II], and 1D). Given that the majority of inositol synthesis in humans occurs in kidney tissue [29], it is reasonable to infer that HEK293T cells would exhibit abundant IP6 expression. This is likely to contribute to enhanced virion maturation efficiency. It has also been suggested that IP6 plays a crucial role in facilitating the assembly of the immature HIV-1 Gag lattice [30]. Consequently, the elevated expression of IP6 in HEK293T cells might enhance the assembly of Gag precursors at the budding site prior to virion release, potentially initiating the trimerization of the MA domain. This, in turn, could enhance the efficient incorporation of Env proteins into progeny virions [31, 32] Another study employing a similar concept of FRET-based assay with different FRET pair proteins revealed that the activation of HIV-1 viral protease occurs during assembly and budding before the release of particles in HEK293T cells [33]. The early maturation observed in HEK293T cells could lead to an increase in progeny likely accounts for the elevated maturation ratio observed in HEK293T cells in our study (Fig. 1). This early maturation could also result in enhanced viral infectivity compared to virions originating from Jurkat cells (Fig. 2)

Several accessory molecules, such as Vif, are not required to produce infectious viruses from HEK293T cells [13, 25, 26]. This is due to the absence of antiviral host factors in HEK293T cells that would normally be counteracted by viral accessory proteins. This discrepancy may hint at the heightened susceptibility of kidney cells to retrovirus infection, potentially attributable to their lack of prior exposure to these viral components. Thus, the environment in HEK293T cells is optimized not only by the absence of antiviral host protein expression but also by the enrichment of host cofactors, which collectively contribute to increased virus infectivity.

Our research addresses a critical gap in the existing literature by presenting a comparative analysis of the efficiency of Gag polyprotein cleavage in virions produced by kidney cells (HEK293T) and T cells (Jurkat). This comparison is vital, considering T cells are the primary source of HIV-1 replication *in vivo*. The observed differences in maturation efficiencies and infectivity rates between HEK293T and Jurkat cells highlight the importance of considering the cellular context in HIV-1 biology and pathogenesis studies. Furthermore, our findings have significant implications for the evaluation of antiviral effects, particularly in the context of long-acting treatments with maturation and capsid inhibitors. These inhibitors target various stages of virus infection, including viral budding and virion maturation [31, 34, 35]. The substantial disparities in viral maturation rates between T-cells and HEK293T cells, the latter of which is a standard cellular resource for *in-vitro* virus production, suggest that relying solely on HEK293T cells could lead to misleading assessments of antiviral efficacy. Therefore, our study indicates the necessity of including T-cell-based assessments to more accurately determine the antiviral efficacy of maturation and capsid inhibitors in virus maturation.

In conclusion, the results of our study not only enhance our understanding of HIV-1 pathogenesis but also emphasize the complexity of the virus life cycle. The unique insights gained into the relationship between virion maturation and infectivity open new pathways for future research, potentially leading to novel therapeutic approaches in the fight against HIV-1.

## Materials and Methods

### Plasmid DNA Construction

The plasmid DNA utilized for HIV-1 Gag-iFRET labeling virus production incorporates an efficient intramolecular FRET pair, ECFPΔC11 and cp173Venus [36]. These genes are positioned between the MA and CA regions of the HIV-1 Gag protein based on the HIV-1 Gag-iGFP construct [37]. The FRET pair genes, along with the junction with the MA and CA regions, are flanked by HIV-1 protease cleavage sites (SQNYPIVQ). This design enables the HIV-1 Protease to specifically cleave at these sites, resulting in a modification of the FRET signal during the maturation of the virion. Additional details are provided in our previous publication [13].

### Virus Production and Cell Culture

HIV-1 Gag-iFRET plasmid DNA was transfected into HEK293T cells using a PEI transfection method (Polysciences) and into Jurkat cells using Lipofectamine LTX transfection reagent (Invitrogen). HEK293T and TZM-bl cells were cultured in Dulbecco’s Modified Eagle’s Medium (Cytiva) supplemented with 10% Fetal Bovine Serum (Gibco), 1% Penicillin-Streptomycin-Glutamine (Invitrogen), and 1% GlutaMax (Gibco)(D10) at 37°C in a 5% CO_2_ environment. Suspended Jurkat cells were cultured in RPMI 1640 media (Cytiva) with the same supplements (R10). To produce FRET-labelled virions, HEK293T cells (7.0 × 10^6^ cells/10 cm dish) and Jurkat cells (10 × 10^6^ cells/10 cm dish) were co-transfected with pHIV-1 Gag-iFRETΔEnv or iFRETΔPRΔEnv, along with the parental plasmids pNL4-3ΔEnv or pNL43ΔPRΔEnv, and the pSVIII-92HT593.1 dual-tropic HIV-1 Env expression plasmid, kindly provided by Dr. Viviana Simon at Mount Sinai, using a transfection ratio of 1:10:2.5. The culture medium was replaced with fresh D10 medium after 3.5 hours for PEI transfection. Virus-containing supernatants were collected 48 hours post-transfection. The supernatants were then filtered through a 0.45 μm sterile polyvinylidene difluoride (PVDF, Millipore) membrane and concentrated up to 10-fold using Lenti-X Concentrator (TaKaRa). The virus pellet was resuspended in 100 μL of Phosphate Buffered Saline (PBS) without phenol red (Cytiva).

### FRET-Based Virion Visualization

To visualize HIV-1 Gag-iFRETΔEnv/iFRETΔPRΔEnv labeled virions, dilute the concentrated virus supernatant 800-fold in 0.22 μm PVDF-filtered Hank’s Balanced Salt Solution (HBSS) without Phenol Red (Cytiva) and load 350 μL onto a glass-bottomed 8-chamber slide (ibidi). The slide was then incubated overnight at 4°C. Single-virion images were captured using an A1R MP+ Multiphoton Confocal Microscope (Nikon). Two sets of 21 images were automatically acquired for each sample under optimal focus conditions. The first set employed a 457.9 nm laser for CFP excitation, capturing emissions through 482 nm/35 nm and 540 nm/30 nm filter cubes for CFP and YFP signals, respectively (FRET images). The second set used a 14.5 nm laser for Venus excitation, with emissions read through a 540 nm/30 nm filter cube to detect the YFP signal. The maturation status was quantified as FRET efficiency relative to signals detected in HIV-1 Gag-iFRETΔ1PRΔEnv labeled virions. Images were captured in RAW ND2 format and converted to TIFF files using Fiji, an image processing package based on ImageJ2. Binary images, generated based on the YFP signal, provided XY coordinates for each particle. The centroid of the XY position for each virion particle was determined, and a circle with a 2-pixel radius centered on the calculated centroid was drawn to represent the virion map. Using this virion map, the FRET signal intensity for each virion was extracted from the raw data, and the FRET ratio (YFP/CFP) was calculated for each particle. Additionally, the mean fluorescent intensity (MFI) of YFP within the circle of the virion map was also extracted from the raw data. Histograms displaying the distribution of FRET ratio values were generated, with 100 bins division, and kernel density estimation curves were plotted alongside. The proportion of the total kernel density estimation area overlapping with the HIV-1 Gag-iFRETΔPRΔEnv area was used to determine the proportion of immature virions. Image data analysis and subsequent calculations of mature and immature virion proportions were performed using an updated in-house MATLAB program [13].

### Infectivity Assays

Viral titers were determined using the HIV-1 Gag p24 DuoSet ELISA (R&D Systems). After seeding TZM-bl cells at 1 × 10^4^ in flat-bottom 96-well plates (CELLTREAT), they were exposed to an equivalent amount of virus (5 ng of HIV-1 p24 total) in the following day. The cells were incubated at 37°C for 48 hours in a CO^2^ incubator.

Subsequently, luciferase activity in the infected cells was measured with the Luciferase Assay System (Promega) using a Molecular Devices SpectraMax Microplate Reader.

### Statistical Analysis

Data were analyzed using standard statistical methods. The significance of differences in virion maturation and infectivity between the two cell types was determined using the Mann-Whitney U test with a *p*-value of less than 0.05, which is considered statistically significant. The statistical differences in YFP MFI between the two cell types were calculated using the Wilcoxon matched-pairs signed-rank test.

## Acknowledgments

We extend our thanks to Dr. Viviana Simon at Mount Sinai for generously providing the pSVIII-92HT593.1 plasmid DNA. In addition, Dr. Luca Sardo at ViiV Healthcare kindly reviewed and provided invaluable comments that enriched this research.

## Author’s contributions

TI conceptualized and designed the study. Experiments were carried out by AM, JC, GH, DM, HT, and TI. The development of the in-house MATLAB script for automated image data analysis was a collaborative effort between KN and TI. Data analysis was conducted by KN and TI. The manuscript was written by TI. TI provides financial support for the research. All authors have made substantial contributions to the work and have given their approval for the final version of the article to be published.

## References

1. Ghosh, A. K., Four decades of continuing innovations in the development of antiretroviral therapy for HIV/AIDS: Progress to date and future challenges. Glob Health Med 2023, 5, (4), 194–198.

2. Landovitz, R. J.; Scott, H.; Deeks, S. G., Prevention, treatment and cure of HIV infection. Nat Rev Microbiol 2023, 21, (10), 657–670.

3. Arts, E. J.; Hazuda, D. J., HIV-1 antiretroviral drug therapy. Cold Spring Harb Perspect Med 2012, 2, (4), a007161.

4. Magden, J.; Kaariainen, L.; Ahola, T., Inhibitors of virus replication: recent developments and prospects. Appl Microbiol Biotechnol 2005, 66, (6), 612–21.

5. Adamson, C. S.; Freed, E. O., Novel approaches to inhibiting HIV-1 replication. Antiviral Res 2010, 85, (1), 119–41.

6. Wiebe, L. I.; Knaus, E. E., Concepts for the design of anti-HIV nucleoside prodrugs for treating cephalic HIV infection. Adv Drug Deliv Rev 1999, 39, (1-3), 63–80.

7. Thenin-Houssier, S.; Valente, S. T., HIV-1 Capsid Inhibitors as Antiretroviral Agents. Curr HIV Res 2016, 14, (3), 270–82.

8. Yant, S. R.; Mulato, A.; Hansen, D.; Tse, W. C.; Niedziela-Majka, A.; Zhang, J. R.; Stepan, G. J.; Jin, D.; Wong, M. H.; Perreira, J. M.; Singer, E.; Papalia, G. A.; Hu, E. Y.; Zheng, J.; Lu, B.; Schroeder, S. D.; Chou, K.; Ahmadyar, S.; Liclican, A.; Yu, H.; Novikov, N.; Paoli, E.; Gonik, D.; Ram, R. R.; Hung, M.; McDougall, W. M.; Brass, A. L.; Sundquist, W. I.; Cihlar, T.; Link, J. O., A highly potent long-acting small-molecule HIV-1 capsid inhibitor with efficacy in a humanized mouse model. Nat Med 2019, 25, (9), 1377–1384.

9. Singh, K.; Gallazzi, F.; Hill, K. J.; Burke, D. H.; Lange, M. J.; Quinn, T. P.; Neogi, U.; Sonnerborg, A., GS-CA Compounds: First-In-Class HIV-1 Capsid Inhibitors Covering Multiple Grounds. Front Microbiol 2019, 10, 1227.

10. Sun, L.; Zhang, X.; Xu, S.; Huang, T.; Song, S.; Cherukupalli, S.; Zhan, P.; Liu, X., An insight on medicinal aspects of novel HIV-1 capsid protein inhibitors. Eur J Med Chem 2021, 217, 113380.

11. Flexner, C.; Owen, A.; Siccardi, M.; Swindells, S., Long-acting drugs and formulations for the treatment and prevention of HIV infection. Int J Antimicrob Agents 2021, 57, (1), 106220.

12. Gillis, E. P.; Parcella, K.; Bowsher, M.; Cook, J. H.; Iwuagwu, C.; Naidu, B. N.; Patel, M.; Peese, K.; Huang, H.; Valera, L.; Wang, C.; Kieltyka, K.; Parker, D. D.; Simmermacher, J.; Arnoult, E.; Nolte, R. T.; Wang, L.; Bender, J. A.; Frennesson, D. B.; Saulnier, M.; Wang, A. X.; Meanwell, N. A.; Belema, M.; Hanumegowda, U.; Jenkins, S.; Krystal, M.; Kadow, J. F.; Cockett, M.; Fridell, R., Potent Long-Acting Inhibitors Targeting the HIV-1 Capsid Based on a Versatile Quinazolin-4-one Scaffold. J Med Chem 2023, 66, (3), 1941–1954.

13. Sarca, A. D.; Sardo, L.; Fukuda, H.; Matsui, H.; Shirakawa, K.; Horikawa, K.; Takaori-Kondo, A.; Izumi, T., FRET-Based Detection and Quantification of HIV-1 Virion Maturation. Front Microbiol 2021, 12, 647452.

14. Chen, B., Molecular Mechanism of HIV-1 Entry. Trends Microbiol 2019, 27, (10), 878–891.

15. Pan, X.; Baldauf, H. M.; Keppler, O. T.; Fackler, O. T., Restrictions to HIV-1 replication in resting CD4+ T lymphocytes. Cell Res 2013, 23, (7), 876–85.

16. Banga, R.; Procopio, F. A.; Noto, A.; Pollakis, G.; Cavassini, M.; Ohmiti, K.; Corpataux, J. M.; de Leval, L.; Pantaleo, G.; Perreau, M., PD-1(+) and follicular helper T cells are responsible for persistent HIV-1 transcription in treated aviremic individuals. Nat Med 2016, 22, (7), 754–61.

17. Berg, R. K.; Rahbek, S. H.; Kofod-Olsen, E.; Holm, C. K.; Melchjorsen, J.; Jensen, D. G.; Hansen, A. L.; Jorgensen, L. B.; Ostergaard, L.; Tolstrup, M.; Larsen, C. S.; Paludan, S. R.; Jakobsen, M. R.; Mogensen, T. H., T cells detect intracellular DNA but fail to induce type I IFN responses: implications for restriction of HIV replication. PLoS One 2014, 9, (1), e84513.

18. Jones, R. B.; Ndhlovu, L. C.; Barbour, J. D.; Sheth, P. M.; Jha, A. R.; Long, B. R.; Wong, J. C.; Satkunarajah, M.; Schweneker, M.; Chapman, J. M.; Gyenes, G.; Vali, B.; Hyrcza, M. D.; Yue, F. Y.; Kovacs, C.; Sassi, A.; Loutfy, M.; Halpenny, R.; Persad, D.; Spotts, G.; Hecht, F. M.; Chun, T. W.; McCune, J. M.; Kaul, R.; Rini, J. M.; Nixon, D. F.; Ostrowski, M. A., Tim-3 expression defines a novel population of dysfunctional T cells with highly elevated frequencies in progressive HIV-1 infection. J Exp Med 2008, 205, (12), 2763–79.

19. Zhang, Z.; Schuler, T.; Zupancic, M.; Wietgrefe, S.; Staskus, K. A.; Reimann, K. A.; Reinhart, T. A.; Rogan, M.; Cavert, W.; Miller, C. J.; Veazey, R. S.; Notermans, D.; Little, S.; Danner, S. A.; Richman, D. D.; Havlir, D.; Wong, J.; Jordan, H. L.; Schacker, T. W.; Racz, P.; Tenner-Racz, K.; Letvin, N. L.; Wolinsky, S.; Haase, A. T., Sexual transmission and propagation of SIV and HIV in resting and activated CD4+ T cells. Science 1999, 286, (5443), 1353–7.

20. Groot, F.; van Capel, T. M.; Schuitemaker, J.; Berkhout, B.; de Jong, E. C., Differential susceptibility of naive, central memory and effector memory T cells to dendritic cell-mediated HIV-1 transmission. Retrovirology 2006, 3, 52.

21. Mattei, S.; Flemming, A.; Anders-Osswein, M.; Krausslich, H. G.; Briggs, J. A.; Muller, B., RNA and Nucleocapsid Are Dispensable for Mature HIV-1 Capsid Assembly. J Virol 2015, 89, (19), 9739–47.

22. Keller, P. W.; Huang, R. K.; England, M. R.; Waki, K.; Cheng, N.; Heymann, J. B.; Craven, R. C.; Freed, E. O.; Steven, A. C., A two-pronged structural analysis of retroviral maturation indicates that core formation proceeds by a disassembly-reassembly pathway rather than a displacive transition. J Virol 2013, 87, (24), 13655–64.

23. Fontana, J.; Jurado, K. A.; Cheng, N.; Ly, N. L.; Fuchs, J. R.; Gorelick, R. J.; Engelman, A. N.; Steven, A. C., Distribution and Redistribution of HIV-1 Nucleocapsid Protein in Immature, Mature, and Integrase-Inhibited Virions: a Role for Integrase in Maturation. J Virol 2015, 89, (19), 9765–80.

24. Hanne, J.; Gottfert, F.; Schimer, J.; Anders-Osswein, M.; Konvalinka, J.; Engelhardt, J.; Muller, B.; Hell, S. W.; Krausslich, H. G., Stimulated Emission Depletion Nanoscopy Reveals Time-Course of Human Immunodeficiency Virus Proteolytic Maturation. ACS Nano 2016, 10, (9), 8215–22.

25. Sheehy, A. M.; Gaddis, N. C.; Choi, J. D.; Malim, M. H., Isolation of a human gene that inhibits HIV-1 infection and is suppressed by the viral Vif protein. Nature 2002, 418, (6898), 646–50.

26. Izumi, T.; Shirakawa, K.; Takaori-Kondo, A., Cytidine deaminases as a weapon against retroviruses and a new target for antiviral therapy. Mini Rev Med Chem 2008, 8, (3), 231–8.

27. Renner, N.; Kleinpeter, A.; Mallery, D. L.; Albecka, A.; Rifat Faysal, K. M.; Bocking, T.; Saiardi, A.; Freed, E. O.; James, L. C., HIV-1 is dependent on its immature lattice to recruit IP6 for mature capsid assembly. Nat Struct Mol Biol 2023, 30, (3), 370–382.

28. Sowd, G. A.; Aiken, C., Correction: Inositol phosphates promote HIV-1 assembly and maturation to facilitate viral spread in human CD4+ T cells. PLoS Pathog 2021, 17, (3), e1009389.

29. Parthasarathy, L. K.; Seelan, R. S.; Tobias, C.; Casanova, M. F.; Parthasarathy, R. N., Mammalian inositol 3-phosphate synthase: its role in the biosynthesis of brain inositol and its clinical use as a psychoactive agent. Subcell Biochem 2006, 39, 293–314.

30. Dick, R. A.; Zadrozny, K. K.; Xu, C.; Schur, F. K. M.; Lyddon, T. D.; Ricana, C. L.; Wagner, J. M.; Perilla, J. R.; Ganser-Pornillos, B. K.; Johnson, M. C.; Pornillos, O.; Vogt, V. M., Inositol phosphates are assembly co-factors for HIV-1. Nature 2018, 560, (7719), 509–512.

31. Freed, E. O., HIV-1 assembly, release and maturation. Nat Rev Microbiol 2015, 13, (8), 484–96.

32. Tedbury, P. R.; Novikova, M.; Alfadhli, A.; Hikichi, Y.; Kagiampakis, I.; KewalRamani, V. N.; Barklis, E.; Freed, E. O., HIV-1 Matrix Trimerization-Impaired Mutants Are Rescued by Matrix Substitutions That Enhance Envelope Glycoprotein Incorporation. J Virol 2019, 94, (1).

33. Tabler, C. O.; Wegman, S. J.; Chen, J.; Shroff, H.; Alhusaini, N.; Tilton, J. C., The HIV-1 Viral Protease Is Activated during Assembly and Budding Prior to Particle Release. J Virol 2022, 96, (9), e0219821.

34. Wang, D.; Lu, W.; Li, F., Pharmacological intervention of HIV-1 maturation. Acta Pharm Sin B 2015, 5, (6), 493–9.

35. Kleinpeter, A. B.; Freed, E. O., HIV-1 Maturation: Lessons Learned from Inhibitors. Viruses 2020, 12, (9).

36. Nagai, T.; Miyawaki, A., A high-throughput method for development of FRET-based indicators for proteolysis. Biochem Biophys Res Commun 2004, 319, (1), 72–7.

37. Hubner, W.; Chen, P.; Del Portillo, A.; Liu, Y.; Gordon, R. E.; Chen, B. K., Sequence of human immunodeficiency virus type 1 (HIV-1) Gag localization and oligomerization monitored with live confocal imaging of a replication-competent, fluorescently tagged HIV-1. J Virol 2007, 81, (22), 12596–607.

